# The Replication-Competent HIV-1 Latent Reservoir is Primarily Established Near the Time of Therapy Initiation

**DOI:** 10.1101/512475

**Authors:** Melissa-Rose Abrahams, Sarah B. Joseph, Nigel Garrett, Lynn Tyers, Matthew Moeser, Nancie Archin, Olivia D. Council, David Matten, Shuntai Zhou, Deelan Doolabh, Colin Anthony, Nilu Goonetilleke, Salim Abdool Karim, David M. Margolis, Sergei Kosakovsky Pond, Carolyn Williamson, Ronald Swanstrom

**Affiliations:** Division of Medical Virology, Institute of Infectious Diseases and Molecular Medicine, University of Cape Town, Cape Town, South Africa.; Department of Microbiology and Immunology, University of North Carolina at Chapel Hill, Chapel Hill, NC, USA.; Lineberger Comprehensive Cancer Center, University of North Carolina at Chapel Hill, Chapel Hill, NC, USA.; Centre for the AIDS Programme of Research in South Africa (CAPRISA), University of KwaZulu-Natal, Durban, South Africa.; UNC HIV Cure Center and Department of Medicine, University of North Carolina at Chapel Hill, Chapel Hill, NC, USA.; Department of Epidemiology, Mailman School of Public Health, Columbia University, New York, NY, USA; Institute for Genomics and Evolutionary Medicine, Temple University, Philadelphia, PA, USA.; National Health Laboratory Services of South Africa.; Department of Biochemistry and Biophysics, University of North Carolina at Chapel Hill, Chapel Hill, NC, USA.

## Abstract

Although antiretroviral therapy (ART) is highly effective at suppressing HIV-1 replication, the virus persists as a latent reservoir in resting CD4+ T cells during therapy. Little is known about the dynamics of reservoir formation and this reservoir forms even when ART is initiated early after infection. The reservoir of individuals who initiate therapy in chronic infection is generally larger and genetically more diverse than that of individuals who initiate in acute infection, suggesting the reservoir is formed continuously throughout untreated infection. To determine when viruses enter the latent reservoir, we compared sequences of replication-competent viruses from resting CD4+ T cells from nine women on therapy to viral sequences circulating in blood collected longitudinally prior to therapy. We found that 78% of viruses from the latent reservoir were most genetically similar to viruses replicating just prior to therapy initiation. This proportion is far greater than expected if the reservoir forms continuously and is always long-lived. Thus, therapy alters the host environment in a way that allows the formation of a majority of the long-lived latent HIV-1 reservoir.

**One Sentence Summary:** Most of the long-lived, replication-competent HIV-1 reservoir is formed at the time of therapy initiation.

## Introduction

Infection with HIV-1 results in active viral replication in the face of the host immune response, eventually leading to the loss of the viral target cell, CD4+ T cells, and immunodeficiency. The use of multiple, potent antiviral drugs stops viral replication and disease progression. However, discontinuation of therapy results in the rapid rebound of virus, indicating that while therapy suppresses viral replication HIV-1 is able to persist in an infectious state for years. The best characterized reservoir for the persistence of HIV-1 during antiretroviral therapy (ART) is as integrated viral DNA in resting CD4+ T cells (*1-3*). One measure of the reservoir is the number of resting CD4+ T cells that can be induced to produce replication-competent virus after stimulation of the cells in culture, called the quantitative virus outgrowth assay (QVOA). Using this assay it has been shown that in people on therapy the latent reservoir has a half-life of 44 months, and approximately one out of a million resting CD4+ T cells can be induced to produce virus (*4, 5*). Given the large number of resting CD4+ T cells in the body, it is impossible to cure HIV-1 by waiting for the infected cells to decay. This problem is exacerbated by the fact that infected cells can clonally expand providing another mechanism for persistence of virus in the body over time (*6-10*).

In people on ART, greater than 90% of the proviral genomes in their resting CD4+ T cells are defective (*11, 12*). These defective genomes may contribute to continued immune activation and exhaustion (*13, 14*), but are not the source of virus in rebound if ART is stopped. In contrast, most intact proviruses are theoretically capable of producing virus, but the frequency of cells harboring intact proviruses is approximately 27 times higher than the frequency of cells that can be induced to produce virus in a QVOA (*12*). In addition, resting CD4+ T cells produce more outgrowth viruses in QVOAs that include extra rounds of cell stimulation (*11*), indicating that the typical QVOA with a single round of stimulation underestimates the number of inducible proviruses. Taken together these results suggest that the reservoir of replication competent proviruses is significantly larger than that measured by standard QVOAs. It is currently unknown whether this discrepancy arises because virus expression from resting CD4+ T-cells is a stochastic process and/or because latency is generated by multiple mechanisms, some of which are not readily reversed in a standard QVOA.

The most widely accepted model of how the reservoir forms involves the infection of a CD4+ T cell as it is transitioning to a resting state. However, little is known about when this happens during the course of infection. The reservoir is present even when ART is initiated early (*15*) (i.e. virus rebounds with the subsequent discontinuation of therapy), and this also suggests that there is early and continuous formation of the reservoir during the period prior to therapy initiation. There are conflicting data on when viral DNA enters the long-lived reservoir with one report claiming there is continuous introduction (*16*) while another report finding that most of the viral DNA in the reservoir comes from virus replicating around the time of therapy initiation (*17*). This is an important question since an understanding of when the reservoir forms could provide new opportunities for limiting its formation, or specifically targeting the reservoir population, as part of a larger cure strategy. In this study we perform the first study analyzing when replication-competent virus enters the long-lived latent reservoir.

We compared sequences of replication-competent viruses induced from on-ART resting CD4+ T cells (outgrowth viruses, OGVs) to viral sequences in blood collected longitudinally pre-ART. We found 78% of OGVs were most similar to virus replicating just prior to therapy initiation. This proportion is far greater than expected if the reservoir forms continuously. Our results indicate that therapy alters the host environment to promote latency and/or dramatically extends the half-life of latently infected cells, suggesting new approaches that could be employed at the time of therapy initiation to block formation of a majority of the long-lived reservoir.

## Results

### Formation of the long-lived viral reservoir in HIV-infected women

We investigated when replication-competent viruses enter the reservoir by examining viruses induced to replicate from resting CD4+ T cells in the context of QVOA. OGVs were isolated from blood-derived cells of nine women on therapy. The women were part of the CAPRISA 002 cohort, based in KwaZulu-Natal, South Africa, who were originally enrolled into the cohort during acute/primary HIV-1 infection (*18*). QVOA measures the replication-competent viral reservoir and estimates the number of resting (CD25-CD69-HLADR-) CD4+ T cells that can produce replication-competent virus following stimulation. This provides a representation of viruses capable of rebound upon ART interruption as opposed to assays that evaluate total viral DNA genomes, the majority of which cannot give rise to replicating virus. The women were ART-naïve for an average of 4.5 years, then initiated ART based on national guidelines in place at the time, and had been on suppressive ART for an average of 4.9 years when blood samples were collected for QVOA (Table 1, Fig. 1). Plasma-derived viral RNA genomes were sequenced at multiple time points pre-ART (on average every six months from acute/early infection) and these evolving sequences were compared to the sequences of the replication-competent OGVs subsequently grown out of the latent reservoir.

**Table 1.** Clinical information for the nine women from the CAPRISA 002 cohort

| Participant ID | *Drug regimen | Time ART naïve (wks) | Time on ART (wks) | CD4 count at ART initiation (cells/μl) | CD4 count at QVOA blood draw (cells/μl) | Number of outgrowth viruses | Number of clonal outgrowth viruses | % Outgrowth viruses from 1 year pre-ART |
| --- | --- | --- | --- | --- | --- | --- | --- | --- |
| CAP188 | EFV/3TC/TDF<br>EFV/FTC/TDF | 247 | 246 | 364 | 430 | 12 | 0 | 67 |
| CAP206 | EFV/3TC/TDF<br>EFV/FTC/TDF | 274 | 287 | 289 | 978 | 21 | 4 | 78 |
| CAP217 | EFV/3TC/TDF | 360 | 238 | 287 | 657 | 17 | 2 | 47 |
| CAP257 | 3TC/NVP/TDF<br>EFV/3TC/TDF<br>EFV/FTC/TDF | 249 | 284 | 170 | 596 | 48 | 8 | 93 |
| CAP287 | EFV/FTC/TDF | 260 | 238 | 411 | 559 | 6 | 0 | 60 |
| CAP288 | EFV/FTC/TDF | 210 | 288 | 318 | 860 | 7 | 0 | 100 |
| CAP302 | EFV/3TC/AZT<br>EFV/3TC/AZT<br>EFV/FTC/TDF | 164 | 269 | 307 | 541 | 8 | 3 | 17 |
| CAP316 | 3TC/AZT/LPV/r<br>EFV/FTC/TDF | 215 | 203 | 436 | 888 | 11 | 0 | 91 |
| CAP336 | EFV/3TC/TDF | 142 | 263 | 164 | 558 | 17 | 8 | 91 |

EFV = Efavirenz, 3TC = Lamivudine, TDF = Tenofovir, NVP = Nevirapine, AZT = Zidovudine, LPV/r
= Lopinavir/Ritonavir,
FTC = Emtricitabine
ART = antiretroviral therapy; QVOA = quantitative virus outgrowth assay
\*Regimen changes indicated below original/prior regimen

**Fig 1.**
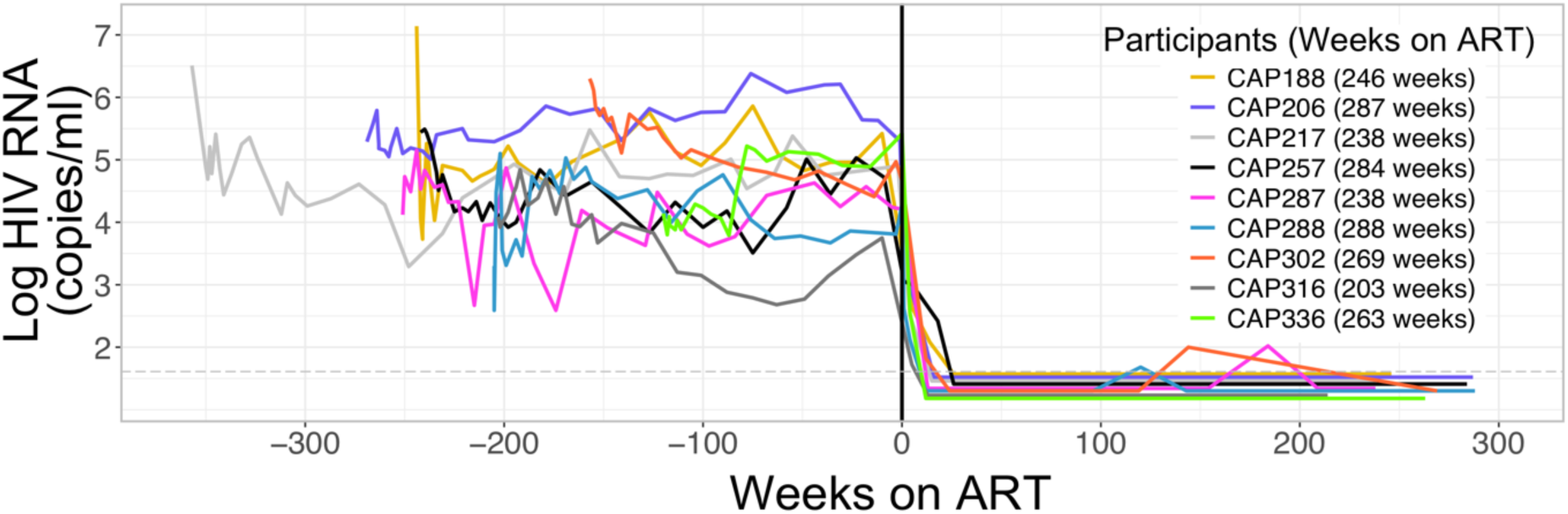
Viral Load and Suppression History of Nine Participants from the CAPRISA 002 Acute Infection Cohort. The graph shows the levels of HIV-1 RNA in the blood pre-and post-therapy, with the time of therapy initiation designated as T=0. Each participant is designated by a different color.

### Sequencing plasma virus from throughout untreated infection and OGVs

We analyzed three viral genes (*env, nef*, and *gag*) known to have rapid rates of evolution driven by immune responses in the host (*19, 20*). Phylogenetic trees of such genomic regions generally adopt a ladder-like structure as the viral population diverges from its homogeneous founder (*21-23*) thereby providing a strong phylogenetic signal for distinguishing populations over time. We estimated when each OGV was seeded into the reservoir by identifying the pre-ART viral sequences that it was most closely related to, and therefore the pre-ART time point when it entered the long-lived reservoir. Phylogenetic placement analysis (*24, 25*) of these three genes supported our estimate of when each OGV entered the reservoir.

An average of nine pre-ART plasma samples were sequenced per participant, using the Illumina MiSeq platform with a Primer ID protocol that allowed us to deeply sample the replicating and evolving viral populations in plasma pre-ART (Supplementary Data Figure 1) while correcting for sequencing errors and template resampling (*26, 27*). Near full-length OGV sequences were generated using the Pacific Biosciences (PacBio) SMRT platform with barcoded primers that likewise allowed for virtually complete error correction. The sequence of each OGV was determined from a bulk PCR product given that the resting CD4+ T cells had been diluted to end point so that a single virus was present in the culture well. Within the different OGVs from a subject, 17% of the sequences were clonal using a definition in which sequences are clonal if they differ by fewer than 5 bases (i.e. 5 differences out of ~ 8845 bases sequenced; Table 1). Since these OGVs were likely derived from cells that had undergone clonal proliferation in the host (*6, 7*), we assumed that clonal OGVs represent a single entry into the reservoir followed by subsequent cellular expansion and therefore included only one representative sequence from each clonal population in summarizing our findings.

### Analysis of the timing of reservoir formation

Phylogenetic analysis revealed that reservoir OGV sequences predominantly clustered with plasma viruses from the year preceding ART initiation (representative trees for two gene regions and three participants are shown in Fig. 2A-C, and trees for all participants are shown in Supplementary Data Figs. 2-10). Overall, a median of 78% of reservoir viruses were most genetically similar to viruses circulating in the year before ART. In four of nine women, over 90% of OGVs, and in seven of nine women at least 60% of OGVs, entered the reservoir in the year proceeding ART (Fig. 3A). In contrast, only 5% of OGVs entered the reservoir during acute/early infection (Table 1). OGVs within an individual inferred to be from time points proximal to ART were more phylogenetically divergent (Fig. 3A, blue sequences) than variants inferred to be from early time points in infection (Fig. 3A, red/orange sequences), further validating our approach to estimating viral entry into the reservoir.

**Fig. 2.**
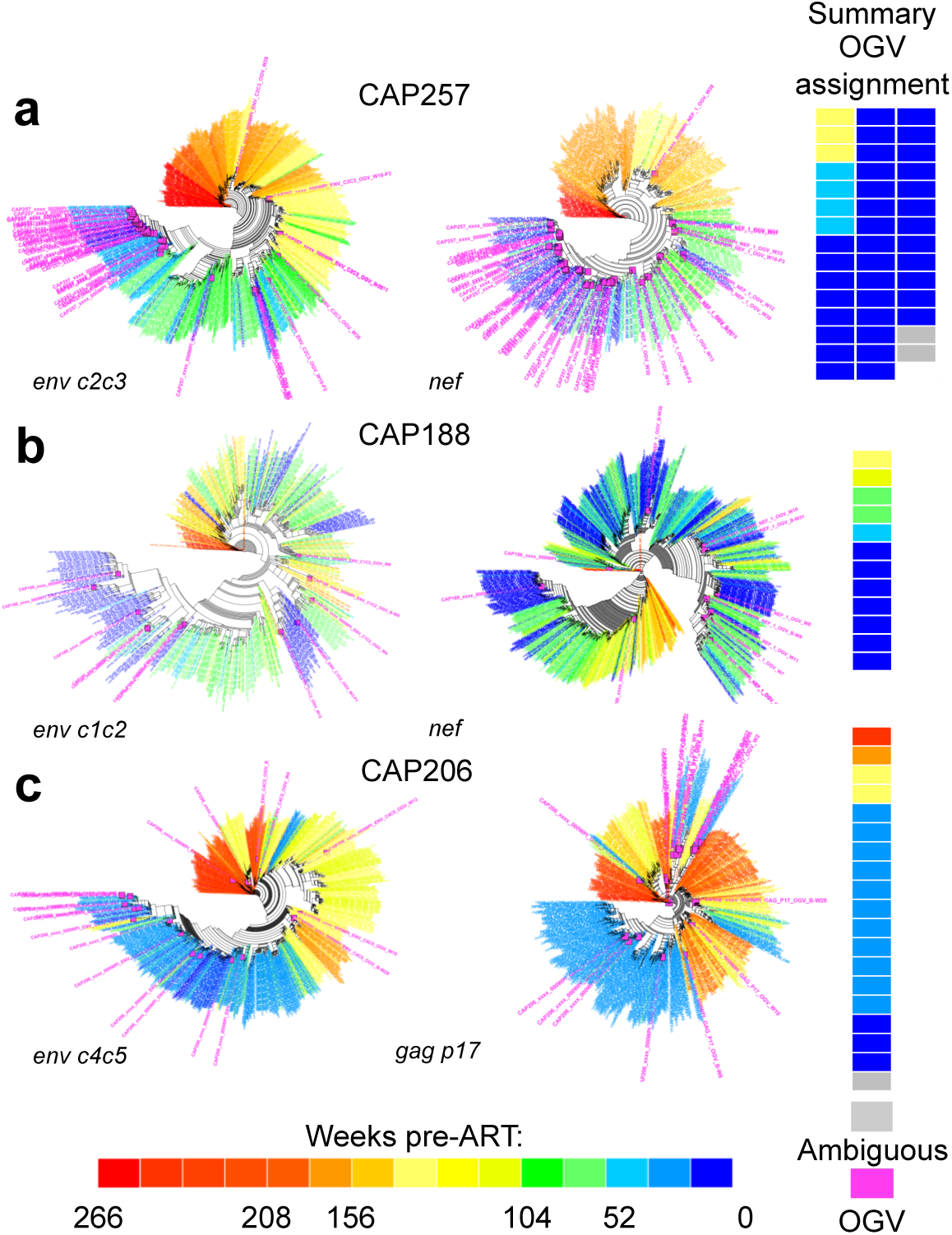
The Majority of Outgrowth Viruses (OGVs) are Most Closely Related to Viruses Replicating Near the Time of ART Initiation. Phylogenetic analyses for two of three genomic regions are shown for three participants. Selected participants each had >10 OGVs with timing assignments for entry into the reservoir in the year preceding ART ranging from 93% to 67% of the OGVs. Heat maps (right) summarize when the closest relatives to each OGV were replicating and therefore when the OGV likely entered the reservoir. Each block represents a genetically unique OGV. (**A)** CAP257 had 44 unique OGVs of which two were designated as ambiguous as they were linked to multiple time points pre-ART. For this individual, 93% of OGVs were most closely related to viruses replicating within one year pre-ART, and 7% to viruses replicating 142 weeks pre-ART. (**B)** CAP188 had 12 unique OGVs, eight (67%) of which were most closely related to viruses replicating within one year pre-ART, and four were most closely related to viruses replicating between 62 and 140 weeks pre-ART. **(C)** CAP206 had 19 unique OGVs and one ambiguous OGV. For this individual, 78% of OGVs were most similar to viruses replicating one year pre-ART, and the remaining were most similar to viruses replicating between 141 weeks and 263 weeks (first year of infection) pre-ART.

**Fig. 3.**
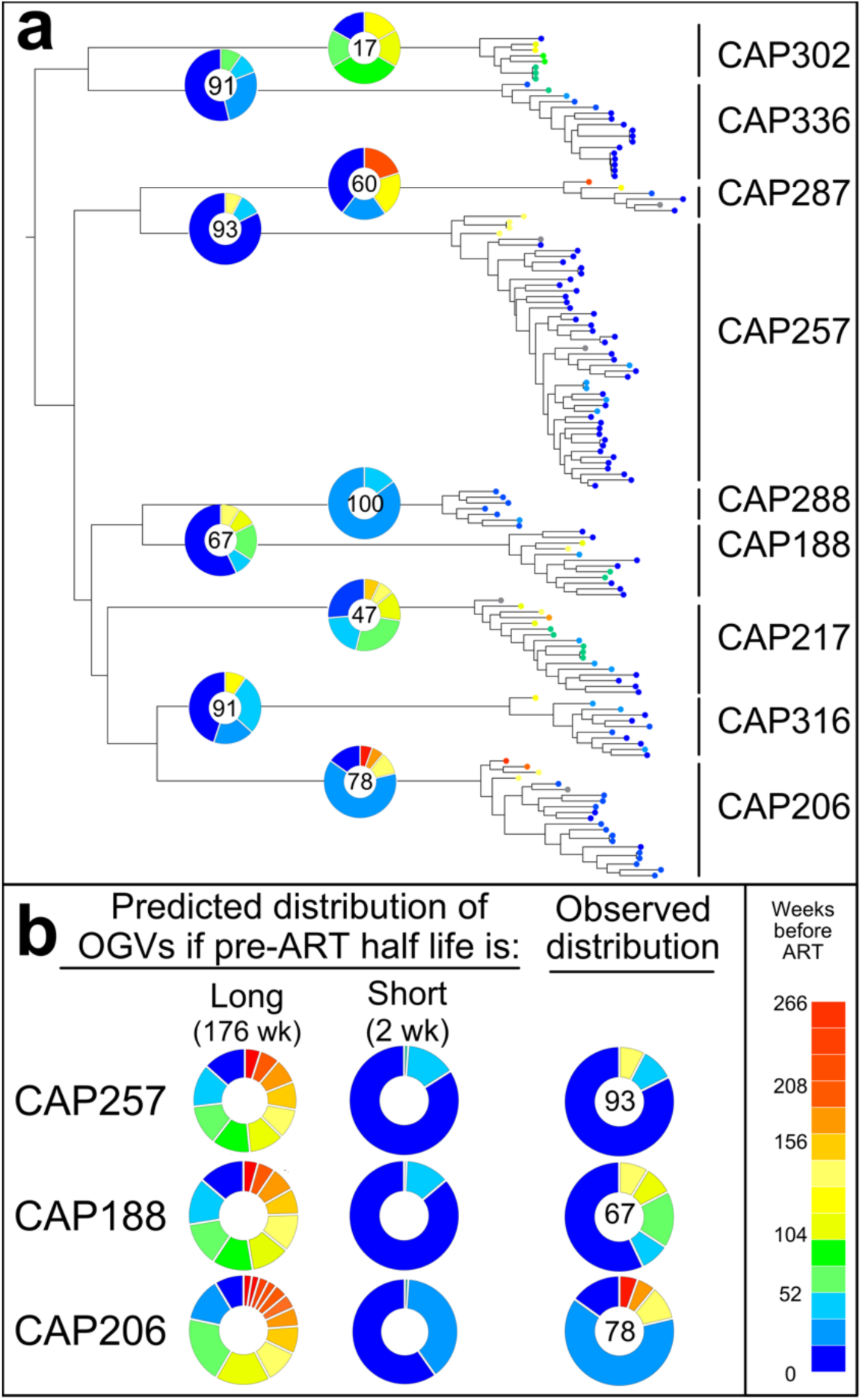
ART-induced changes in cellular half-life of latently infected cells may explain why cells infected near the time of ART initiation comprise the majority of the inducible replication-competent reservoir. **(A)** All outgrowth virus (OGV) sequences are shown in an approximate maximum-likelihood tree. Branch tips are colored according to the estimated time when each OGV entered the reservoir. For each participant, the OGV timing distribution is illustrated in a pie chart with the percentage of OGVs from the year pre-ART listed. Overall, 78% of OGVs were produced by cells infected near the time of ART initiation. (**B)** Two models were considered to explain this pattern. One model assumes that latently infected cells have a long half-life of 176 weeks (44 months) both prior to and during therapy, and predicts that the reservoir will contain variants from throughout the untreated infection. The other model assumes that latently infected cells have a short half-life in untreated infection (here chosen as 2 weeks) that then stabilizes to a long half-life during therapy. For the majority of participants (three shown here), the observed data most closely resembles the predicted pattern if latently infected cells have a short half-life prior to ART initiation and then decay slowly after ART initiation.

### Modeling formation of the long-lived HIV-1 reservoir

Our observations indicate that cells infected with HIV-1 near the time of ART initiation (or clones of those cells) are more likely to contribute to the long-lived, replication-competent reservoir than cells infected at earlier time points. We considered two simple models to explain our findings. The first model assumes that infected resting CD4+ T cells, which have been observed in untreated infection (*28*), have a long half-life of 44 months (176 weeks) comparable to the half-life on therapy (Fig. 3B, left panel) (*4, 5*). Under this model, the reservoir will contain variants sampled throughout untreated infection with those from later in infection being slightly more common. The second model assumes that infected resting CD4+ T cells have a short half-life in untreated HIV-1 infection (here chosen as 2 weeks) that then stabilizes to a longer half-life with or shortly after the onset of effective ART (Fig. 3B, middle panel). Similar to our observations (Fig. 3B, right panel), this second model predicts that most of the reservoir will be seeded by viruses replicating around the time of ART initiation. Thus, our observed high frequency of viruses replicating proximal to ART initiation in the latent reservoir can be explained by infected resting CD4+ T cells from early in infection rapidly decaying before they contribute to the long-lived reservoir.

## Discussion

In this study we have taken advantage of the availability of archived pre-therapy viremic samples in the CAPRISA 002 cohort to establish a detailed molecular clock of viral evolution in each of 9 women enrolled from the time of primary infection. We were able to use this clock to determine when viruses entered the long-lived latent reservoir that had persisted after over 4 years on therapy. In most women the majority of the viruses entered the reservoir around the time of therapy initiation, suggesting that it is the initiation of ART that is indirectly changing the host environment by suppressing viral replication to now favor the establishment of long-lived cells, some of which are latently infected with HIV-1.

While we have studied the timing of entry of the replication competent portion of the reservoir, Brodin *et al.* (*17*) and Jones *et al.* (*16*) recently characterized the formation of the long-lived proviral DNA reservoir, which is predominantly defective (*11, 12*). Our findings representing the replication-competent reservoir are consistent with the observations of Brodin *et al*. (*17*) who examined the viral DNA reservoir in 10 individuals and found that it was skewed toward variants present in the year preceding ART, although to a lesser extent than identified here. The results of Brodin *et al*. differ from those of Jones *et al*. who inferred diversity in the viral DNA reservoir that was reflective of viral evolution over the entire course of infection in two individuals. However, their analysis was based on sequences from a limited number of pre-ART time points (all during chronic infection) and thus required modeling to extrapolate a molecular clock. In contrast, we were able to sequence virus from acute/early infection onward to the start of therapy and estimate that only a small fraction of the sampled OGV populations were seeded prior to the initiation of therapy, and in 8/9 participants there was no evidence of continuous formation. In only one participant (CAP302) was the replication-competent reservoir primarily seeded from early to mid-stages of the untreated infection (Fig. 3A, Extended Data Figure 8). Overall our data and modeling analysis suggest that much of the long-lived HIV-1 reservoir forms following the initiation of ART which also coincides with rapid declines in HIV-1 viral load, increases in CD4+ T cell counts, and reductions in markers of immune activation.

Untreated HIV-1 infection is characterized by increased immune activation and dysfunction, which are significantly reversed after ART initiation. Most relevant to this study is the loss of CD127+(IL-7 receptor) memory CD4+ T cells during untreated infection (*29-31*). IL-7 signaling through the CD127 receptor is essential for the transition of effector to memory CD4+ T cells as well as the subsequent maintenance of memory cells (*32*). With ART initiation, the number of CD127+ memory CD4+ T cells increases (*29, 30, 33*), most likely due to reductions in inflammation (*34*). These results suggest a model in which the increase in CD127 expression during ART allows latently infected cells to become long-lived as they now become responsive to IL-7 (Fig. 4). Consistent with this model in which much of the reservoir should be in cells sensitive to IL-7, administering IL-7 to HIV-infected individuals on ART increases the size of the HIV-1 reservoir (*35*) and the rate of reservoir decay is slower when IL-7 is higher (*36*).

**Fig 4.**
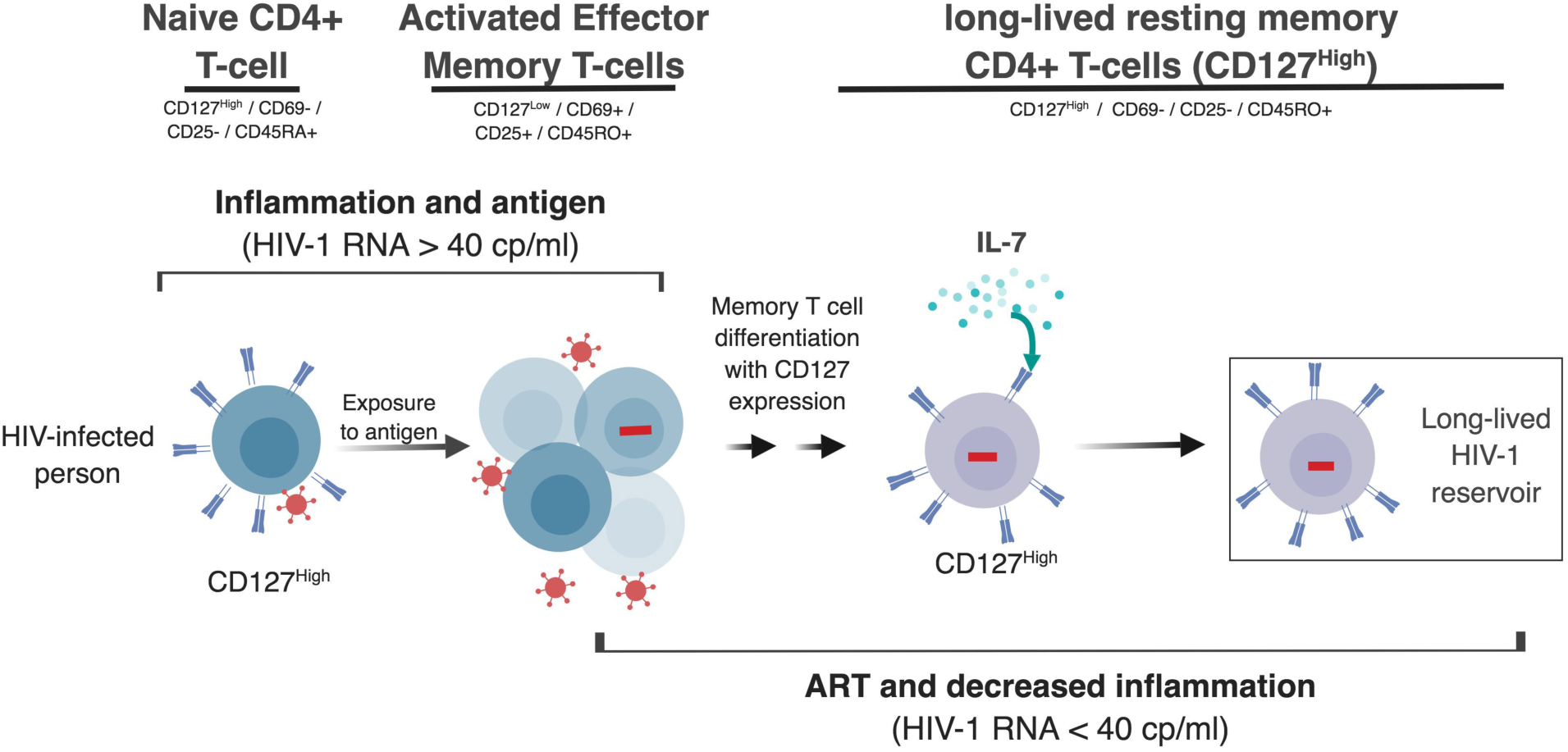
Model of the Relationship Between Viral Suppression and Formation of the Long-lived HIV-1 Reservoir. IL-7 signaling through the CD127 receptor regulates memory CD4+ T cell differentiation, survival and proliferation. Immune activation during untreated HIV-1 infection reduces CD127 expression, thus limiting the transition from effector to differentiated memory cell, and the same signaling may antagonize viral silencing. Viral suppression by ART reduces inflammation and increases CD127 expression. Thus, prior to ART HIV-infected cells are unlikely to be long-lived. In contrast, reductions in inflammation after ART initiation may allow HIV-infected cells to increased CD127 expression and develop the long-lived phenotype that characterizes the HIV-1 reservoir.

These results suggest that therapies that limit CD4+ T cell memory generation, perhaps through limiting IL-7/IL-7R signaling in the window from the time of ART initiation to when all viral replication is fully suppressed, could significantly limit reservoir formation. Such an approach would not eliminate all of the reservoir, as it is clear that latency can be seeded at different points during untreated infection. However, blocking the formation of a majority of the reservoir at the time of therapy initiation would greatly reduce the size of the reservoir, lowering the barrier to eradicating HIV-1 from an infected host.

## Materials and Methods

### Study participants

The CAPRISA 002 cohort is comprised of women from rural and urban KwaZulu-Natal, South Africa, who were identified during their acute infection with subtype C HIV-1 and followed longitudinally. Participants in the cohort gave blood samples every 3-6 months and were provided antiretroviral therapy (ART) based on the prevailing in-country guidelines. The women were retained in the cohort for up to five years after therapy initiation. We identified nine women in the cohort who had been ART naïve for at least three years prior to initiating therapy, and who had been on ART for a minimum of 4 years. The women provided a 200 ml blood draw while on therapy from which peripheral blood mononuclear cells (PBMCs) were isolated and cryopreserved in liquid nitrogen. Women were excluded based on pregnancy and hemoglobin levels lower than 10g/dL. The use of stored samples and the collection and analysis of new samples were done under a protocol approved by the Biomedical Research Ethics Committee of the University of KwaZulu Natal (BE178/150) and the Human Research Ethics Committee of the University of Cape Town (588/2015) in South Africa, and at the University of North Carolina in the United States.

### Viral outgrowth assay

Resting CD4+ T cells were isolated from PBMCs by negative selection with CD25 depletion (Custom kit, Stem Cell). Purified cells were then examined by flow cytometry (LSRII Fortessa) for expression of CD69 (PE, BD), CD25 (APC, BD), CD8 (FITC, BD), CD4 (PerCP-Cy5.5, BD) and viability (aqua live/dead, ThermoFisher). Data were analysed by FlowJo (version 10.4.2).

Resting cells were cultured at 100,000 cells/well with 1.5 µg/ml of highly purified phytohemagglutinin (PHA, Remel-PHA, Thermo Scientific), 60 U/ml of interleukin 2, and irradiated PBMCs from a seronegative donor, and virus outgrowth was facilitated as previously described (*4*). On days 15 and 19, cultures were tested for viral p24 capsid protein production using an enzyme-linked immunosorbent assay. Culture supernatants positive for p24 were stored at −80ºC.

### Sequencing of viral populations in blood plasma pre-ART

Viral RNA copies were extracted from blood plasma using the QIAamp viral RNA Mini kit (Qiagen). The purified RNA was reverse transcribed to cDNA using Superscript III/IV Reverse Transcriptase (Invitrogen). We used the Primer ID method (*26*) which tags each RNA template through its cDNA primer with a unique 12 nucleotide long identifier. This allows amplified sequences to be grouped according to their Primer ID tags, from which a consensus sequence can then be derived for each individual template. Multiple cDNA primers were used during the cDNA synthesis step for each sample to generate cDNAs corresponding to multiple regions of the genome and allow multiplexing of the PCR amplicons followed by sequencing. The gene-specific sequence of the cDNA primer was used as an index for the specific region of the genome. Two separate cDNA reactions were performed for each sample to allow efficient use of the available RNA to sequence most of the viral genome. The cDNA products were purified twice using Agencourt RNA Clean XP magnetic beads (Beckman Coulter). Multiplexing during PCR was accomplished by including a common PCR primer binding sequence at the 5’ end of each cDNA primer and a gene-specific forward primer for each amplicon/region. PCR amplification was performed using the KAPA2G Fast Multiplex Mix (Kapa Biosystems) and an equal molar amount of each forward primer and an excess of the universal reverse primer. A second PCR step using the Expand High Fidelity PCR System (Roche) allowed the incorporation of the Illumina MiSeq version 2 indexes and adapters. PCR products were purified using SPRIselect beads (Beckman Coulter), with a 0.6:1 μl ratio of beads to PCR product (to allow for size exclusion of residual primers) following both PCR steps. The final purified amplicon libraries were quantified using the Qubit dsDNA HS assay (Invitrogen, CA), pooled at equimolar ratios and purified again using SPRIselect beads. Illumina MiSeq 2×300 base paired-end sequencing was performed on the multiplexed amplicons.

### Sequencing of outgrowth virus populations

Viral RNA was isolated from p24-positive wells and converted to cDNA using Superscript III Reverse Transcriptase and an oligo(dT) primer. The 3’ and 5’ half genomes were amplified in separate PCR reactions using barcoded primers, and the PCR products were gel purified. The SMARTbell Template Prep Kit (PacBio) was used to add adaptors to amplicons, and amplicons were then submitted for PacBio sequencing (movie time of 10 hours). Sequences were grouped by barcode and high-quality sequences were analyzed using the PacBio Long Amplicon Analysis (LAA) package. The 3’ and 5’ amplicons for the same virus were joined and visually screened to confirm that open reading frames were intact.

### Illumina MiSeq data processing

Raw reads were processed using a custom pipeline written in Python and R programming languages. The MotifBinner2.R program (https://github.com/HIVDiversity/MotifBinner2) carries out quality filtering, merging of overlapping paired-end reads and implements the Primer ID processing method described by Zhou et al (*27*). The resulting Primer ID consensus sequences were processed with an in-house pipeline (https://github.com/ColinAnthony/NGS_processing_pipeline) which removes sequences with degenerate bases or deletions >50 base pairs long, filters out any contaminant non-target gene sequences, and codon aligns the cleaned longitudinal sequences for each gene region. This was achieved by aligning the translated sequences using MAFFT (*37*) and subsequently back-translating the alignment. Alignments were visually inspected to confirm alignment accuracy. Columns in the alignment with > 90% gap characters and (where applicable) hyper-variable regions of the *env* gene that could not be reliably aligned were removed. The variant detection level (at 95% confidence) for each dataset was calculated using the following binomial equation:

Variant frequency (95% probability) = 1 - (0.05 * (1 / sequencing depth)).

Finally, for each participant, sequences from each time point and for each gene region were collapsed to a single sequence for each group of template consensus sequences with 100% identity.

### Phylogenetic and statistical analyses

Phylogenetic trees were generated using two methods: (i) Approximately-Maximum-Likelihood trees were constructed using FastTree (*38*), using a general time reversible model of evolution, and (ii) by doing amino acid alignments in MUSCLE, constructing trees using FastTree and performing phylogenetic placement of OGVs using in-house software. The most closely related pre-ART sequence to each of the outgrowth virus sequences was determined by both automated leaf analysis using the python library ETE3 (*39*) and visual inspection. We performed phylogenetic placement (*24, 25*) of each individual outgrowth virus on the reference pre-ART viral RNA phylogeny. This methodology has been successful in the statistically rigorous classification of sequences of unknown origin in metagenomics (*40*), immunomics, and viral diversity classification (*41*). For each DNA sequence (S) and pre-therapy time-point (T), phylogenetic placement obtains a score between 0 and 1, representing how certain we are that sequence S originated from time-point T.

## Supporting information

Supplemental Material for Abrahams et al.

## Supplementary Materials

Figures

Fig. S1. Longitudinal MiSeq Sampling Depth for Nine Women from the CAPRISA 002 Cohort.

Fig. S2. Timing of Reservoir Outgrowth Viruses for Participant CAP188.

Fig. S3. Timing of Reservoir Outgrowth Viruses for Participant CAP206.

Fig. S4. Timing of Reservoir Outgrowth Viruses for Participant CAP217.

Fig. S5. Timing of Reservoir Outgrowth Viruses for Participant CAP257.

Fig. S6. Timing of Reservoir Outgrowth Viruses for Participant CAP287.

Fig. S7. Timing of Reservoir Outgrowth Viruses for Participant CAP288.

Fig. S8. Timing of Reservoir Outgrowth Viruses for Participant CAP302.

Fig. S9. Timing of Reservoir Outgrowth Viruses for Participant CAP316.

Fig. S10. Timing of Reservoir Outgrowth Viruses for Participant CAP336.

## Acknowledgments

We would like to acknowledge all participants of the CAPRISA 002 acute infection cohort as well as the staff at the Vulindlela and eThekwini Clinical Research Sites, KwaZulu-Natal, South Africa.

## Author contributions

CW and RS proposed, designed and supervised this study. M-RA, SBJ, and NGarrett directed all of the data collection, experiments and data analyses. MM, NA, OC, LT, SZ, DD, performed all of the experiments. Phylogenetic analyses were performed by SKP, CA, MM, and DM. The manuscript was written by M-RA, SBJ, N Goonetilleke. CW, and RS with editorial help from NGarrett, SAK, DMM, and SKP.

## Data and materials availability

All sequences will be deposited in public databases and accession numbers provided prior to publication. The near full-length genome sequences will be deposited in GenBank. The MiSeq sequences will be deposited in the Sequencing Read Archive.

