## Supplemental Material for Abrahams et al. for "The Replication-Competent HIV-1 Latent Reservoir is Primarily Established Near the Time of Therapy Initiation"

**Supplemental Data**

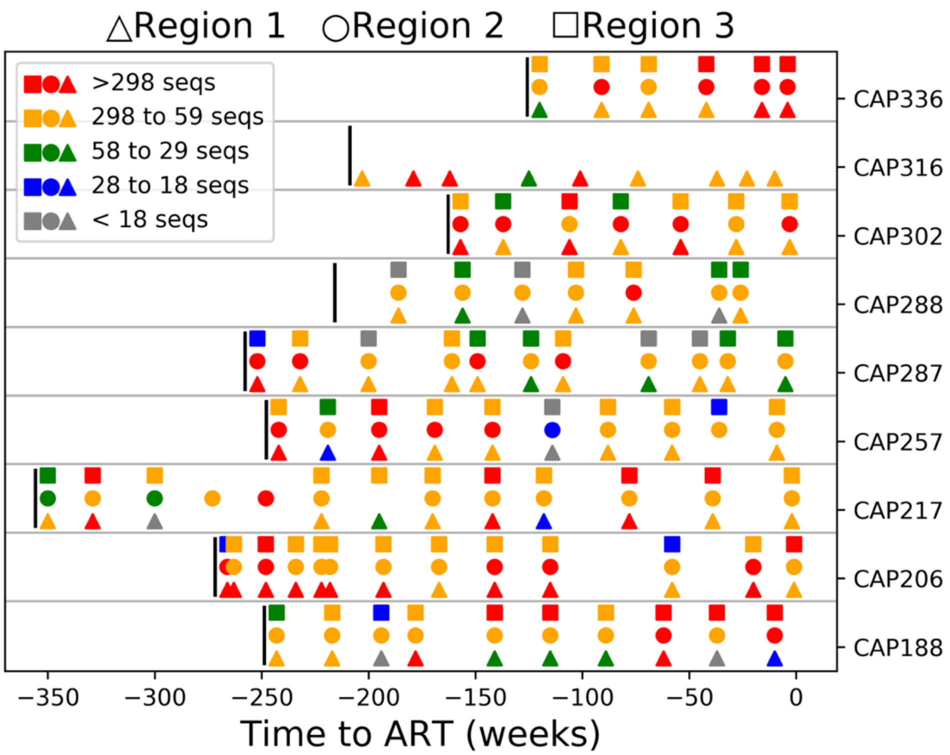


**Fig. S1. Longitudinal MiSeq Sampling Depth for Nine Women from the** CAPRISA 002 Cohort. The sequencing depth (PID-consensus sequences) is indicated over the sampling period for each participant for each gene region analyzed. The enrollment time point for each individual is indicated by a vertical bar. Sequencing depth is indicated by color (red, yellow, green, blue and grey), to indicate 95% probability variant detection limits of <1%, 1-5%, 5-10%, 10-15% and >15%, respectively. Region 1 (triangle), 2 (circle) and 3 (square) correspond to the following regions for participant CAP188: *env c1c2*, *env c2c3*, and *nef*; CAP206: *env c1c2*, *env v4c5*, and *gag p17*; CAP217: *env c1c2*, *env c2c3*, and *env v4c5*; CAP257: *env c1c2*, *env c2c3*, and *nef*; CAP287: *env c2c3*, *env v4c5*, and *gag p17*; CAP288: *env c1c2*, *env c2c3*, and *gag p17*; CAP302: *env c2c3*, *env v4c5*, and *nef*; CAP316: *env c2c3*; CAP336: *env c1c2*, *env v4c5*, and *nef.*

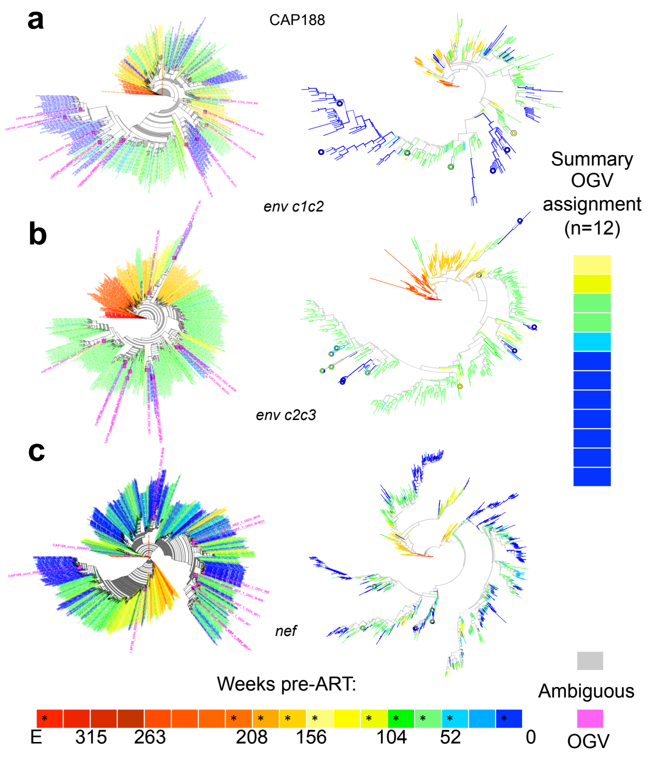


**Fig. S2. Timing of Reservoir Outgrowth Viruses for Participant CAP188.** Two applications of Approximately Maximum-Likelihood trees were used for each of the three gene regions. In one application (on the left) the OGV sequences were included in the alignment and are shown in pink. In the other application phylogenic placement was used to place the OGV sequences in the tree (on the right) to assess the probability of placement illustrated in colored doughnuts for **a,** *env c1c2*, **b,** *env c2c3*, and **c,** *nef*. Sequence labels (left tree panel) and branches/doughnuts (right tree panel) are colored according to time pre-treatment as indicated in the color key. Sampling times are indicated by asterisks on the color key. The summarized time assignment for each unique outgrowth virus is indicated in the heat map on the far right. Colors in the heat map correspond to those in the trees and color key. E = acute infection study enrollment time point.

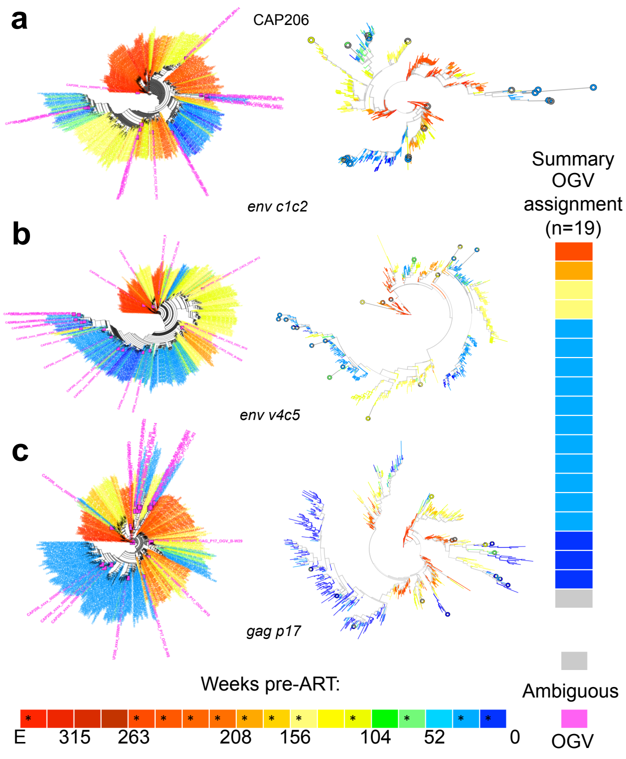


**Fig. S3. Timing of Reservoir Outgrowth Viruses for Participant CAP206.** Two applications of Approximately Maximum-Likelihood trees were used for each of the three gene regions. In one application (on the left) the OGV sequences were included in the alignment and are shown in pink. In the other application phylogenic placement was used to place the OGV sequences in the tree (on the right) to assess the probability of placement illustrated in colored doughnuts for **a,** *env c1c2*, **b,** *env v4c5*, and **c,** *gag p17*. Sequence labels (left tree panel) and branches/doughnuts (right tree panel) are colored according to time pre-treatment as indicated in the color key. Sampling times are indicated by asterisks on the color key. The summarized time assignment for each unique outgrowth virus is indicated in the heat map on the far right. Colors in the heat map correspond to those in the trees and color key. E = acute infection study enrollment time point.

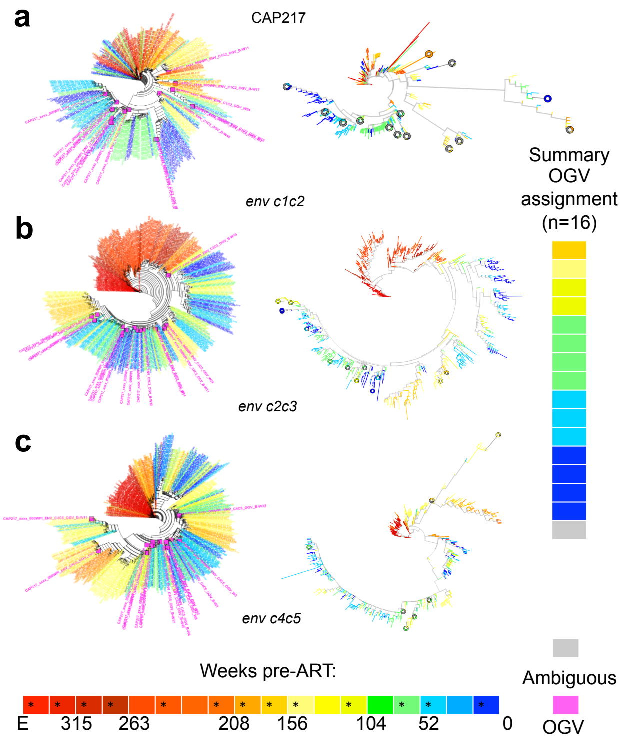


**Fig. S4. Timing of Reservoir Outgrowth Viruses for Participant CAP217.** Two applications of Approximately Maximum-Likelihood trees were used for each of the three gene regions. In one application (on the left) the OGV sequences were included in the alignment and are shown in pink. In the other application phylogenic placement was used to place the OGV sequences in the tree (on the right) to assess the probability of placement illustrated in colored doughnuts for **a,** *env c1c2*, **b,** *env c2c3*, and **c,** *env v4c5*. Sequence labels (left tree panel) and branches/doughnuts (right tree panel) are colored according to time pre-treatment as indicated in the color key. Sampling times are indicated by asterisks on the color key. The summarized time assignment for each unique outgrowth virus is indicated in the heat map on the far right. Colors in the heat map correspond to those in the trees and color key. E = acute infection study enrollment time point.

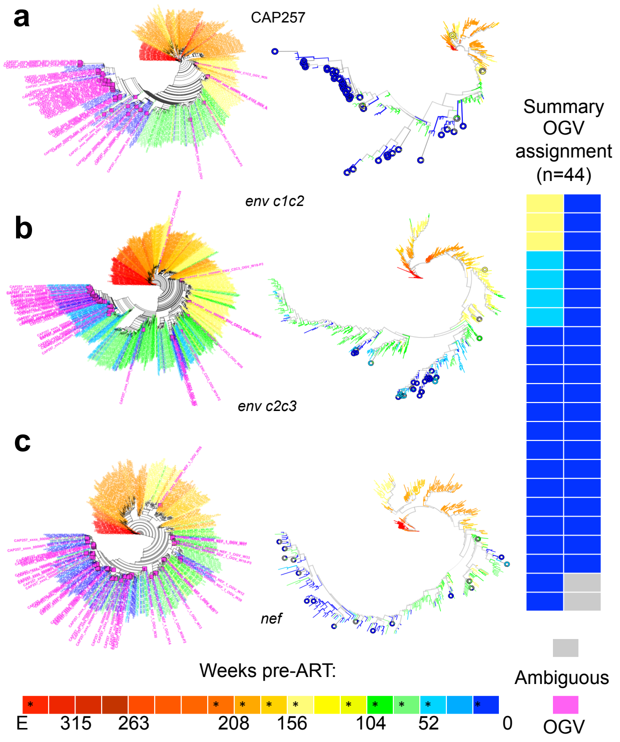


**Fig. S5. Timing of Reservoir Outgrowth Viruses for Participant CAP257.** Two applications of Approximately Maximum-Likelihood trees were used for each of the three gene regions. In one application (on the left) the OGV sequences were included in the alignment and are shown in pink. In the other application phylogenic placement was used to place the OGV sequences in the tree (on the right) to assess the probability of placement illustrated in colored doughnuts for **a,** *env c1c2*, **b,** *env c2c3*, and **c,** *nef*. Sequence labels (left tree panel) and branches/doughnuts (right tree panel) are colored according to time pre-treatment as indicated in the color key. Sampling times are indicated by asterisks on the color key. The summarized time assignment for each unique outgrowth virus is indicated in the heat map on the far right. Colors in the heat map correspond to those in the trees and color key. E = acute infection study enrollment time point.

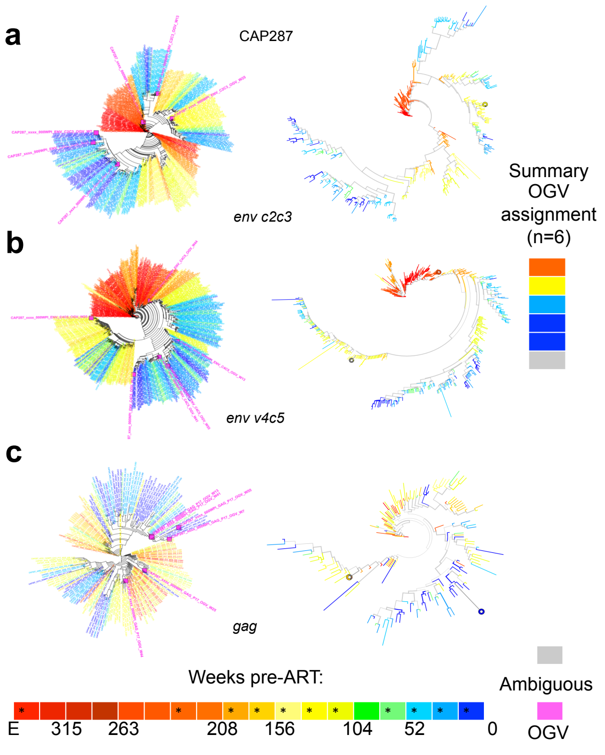


**Fig. S6. Timing of Reservoir Outgrowth Viruses for Participant CAP287.** Two applications of Approximately Maximum-Likelihood trees were used for each of the three gene regions. In one application (on the left) the OGV sequences were included in the alignment and are shown in pink. In the other application phylogenic placement was used to place the OGV sequences in the tree (on the right) to assess the probability of placement illustrated in colored doughnuts for **a,** *env c2c3*, **b,** *env v4c5*, and **c,** *gag p17*. Sequence labels (left tree panel) and branches/doughnuts (right tree panel) are colored according to time pre-treatment as indicated in the color key. Sampling times are indicated by asterisks on the color key. The summarized time assignment for each unique outgrowth virus is indicated in the heat map on the far right. Colors in the heat map correspond to those in the trees and color key. E = acute infection study enrollment time point.

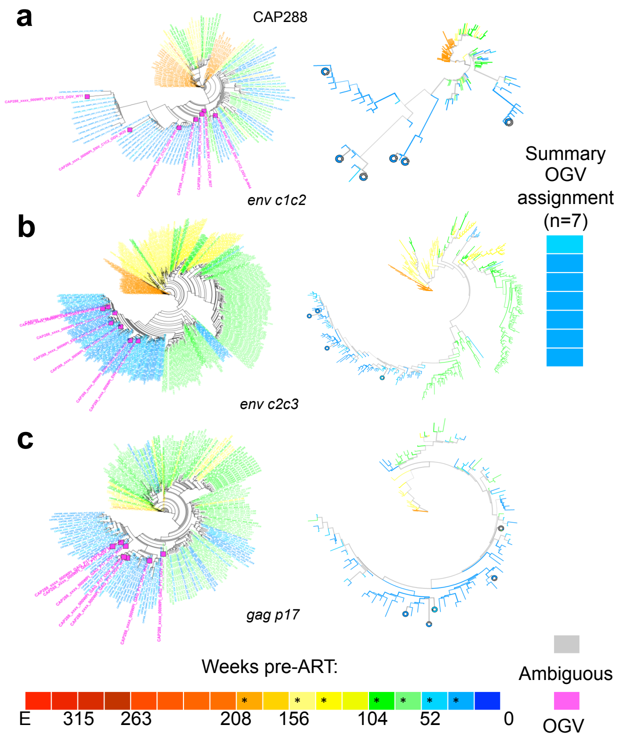


**Fig. S7. Timing of Reservoir Outgrowth Viruses for Participant CAP288.** Two applications of Approximately Maximum-Likelihood trees were used for each of the three gene regions. In one application (on the left) the OGV sequences were included in the alignment and are shown in pink. In the other application phylogenic placement was used to place the OGV sequences in the tree (on the right) to assess the probability of placement illustrated in colored doughnuts for **a,** *env c1c2*, **b,** *env c2c3*, and **c,** *gag p17*. Sequence labels (left tree panel) and branches/doughnuts (right tree panel) are colored according to time pre-treatment as indicated in the color key. Sampling times are indicated by asterisks on the color key. The summarized time assignment for each unique outgrowth virus is indicated in the heat map on the far right. Colors in the heat map correspond to those in the trees and color key. E = acute infection study enrollment time point.

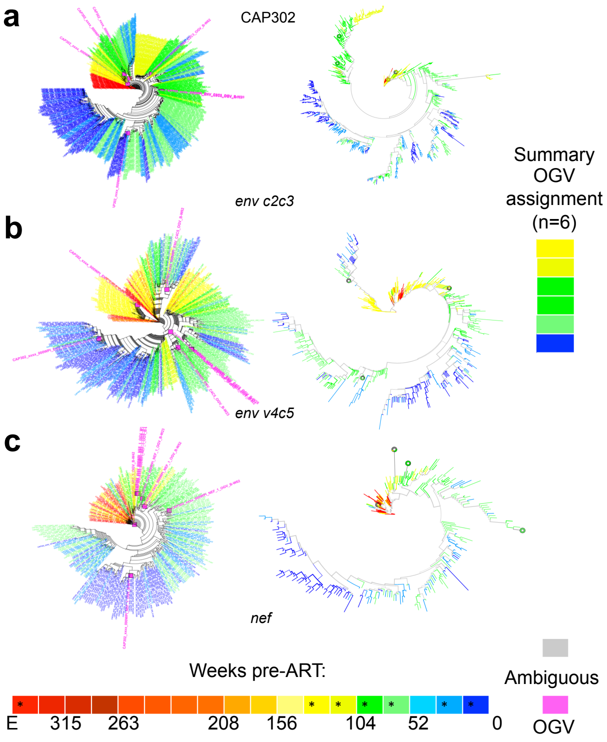


**Fig. S8. Timing of Reservoir Outgrowth Viruses for Participant CAP302.** Two applications of Approximately Maximum-Likelihood trees were used for each of the three gene regions. In one application (on the left) the OGV sequences were included in the alignment and are shown in pink. In the other application phylogenic placement was used to place the OGV sequences in the tree (on the right) to assess the probability of placement illustrated in colored doughnuts for **a,** *env c2c3*, **b,** *env v4c5*, and **c,** *nef*. Sequence labels (left tree panel) and branches/doughnuts (right tree panel) are colored according to time pre-treatment as indicated in the color key. Sampling times are indicated by asterisks on the color key. The summarized time assignment for each unique outgrowth virus is indicated in the heat map on the far right. Colors in the heat map correspond to those in the trees and color key. E = acute infection study enrollment time point.

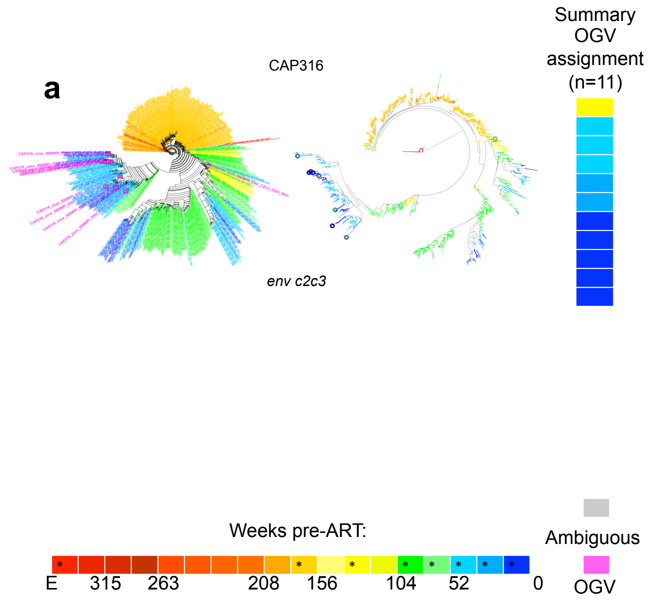


**Fig. S9. Timing of Reservoir Outgrowth Viruses for Participant CAP316.** Two applications of Approximately Maximum-Likelihood trees were used for the gene region shown. In one application (on the left) the OGV sequences were included in the alignment and are shown in pink. In the other application phylogenic placement was used to place the OGV sequences in the tree (on the right) to assess the probability of placement illustrated in colored doughnuts for *env c2c3*. Sequence labels (left tree panel) and branches/doughnuts (right tree panel) are colored according to time pre-treatment as indicated in the color key. Sampling times are indicated by asterisks on the color key. The summarized time assignment for each unique outgrowth virus is indicated in the heat map on the far right. Colors in the heat map correspond to those in the trees and color key. E = acute infection study enrollment time point.

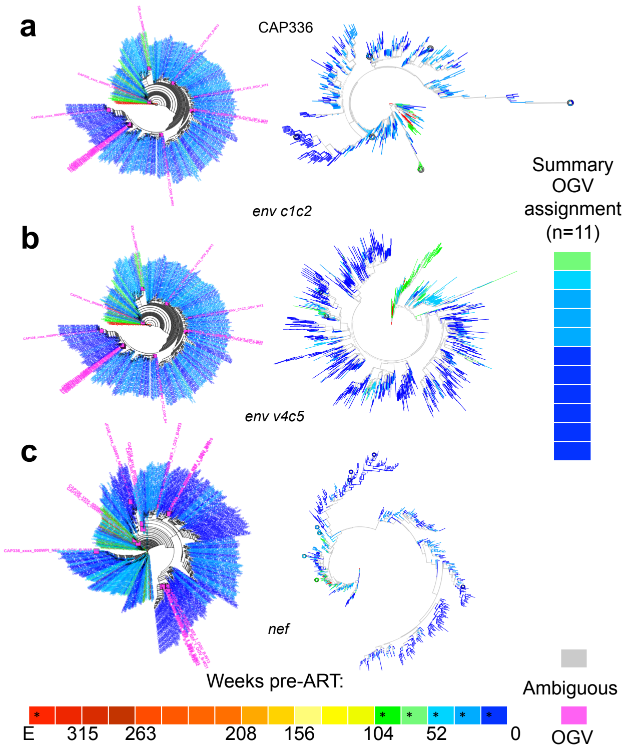


**Fig. S10. Timing of Reservoir Outgrowth Viruses for Participant CAP336.** Two applications of Approximately Maximum-Likelihood trees were used for each of the three gene regions. In one application (on the left) the OGV sequences were included in the alignment and are shown in pink. In the other application phylogenic placement was used to place the OGV sequences in the tree (on the right) to assess the probability of placement illustrated in colored doughnuts for **a,** *env c1c2*, **b,** *env v4c5*, and **c,** *nef*. Sequence labels (left tree panel) and branches/doughnuts (right tree panel) are colored according to time pre-treatment as indicated in the color key. Sampling times are indicated by asterisks on the color key. The summarized time assignment for each unique outgrowth virus is indicated in the heat map on the far right. Colors in the heat map correspond to those in the trees and color key. E = acute infection study enrollment time point.
